# A secreted *Echinococcus multilocularis* activin A homologue promotes regulatory T cell expansion

**DOI:** 10.1101/618140

**Authors:** Justin Komguep Nono, Manfred B. Lutz, Klaus Brehm

**Author notes:** Corresponding authors (KB); (MBL).

## Abstract

**Background:** Alveolar echinococcosis (AE), caused by the metacestode larval stage of the fox-tapeworm *Echinococcus multilocularis*, is a chronic zoonosis associated with significant modulation of the host immune response. A role of regulatory T-cells (Treg) in generating an immunosuppressive environment around the metacestode during chronic disease has been reported, but the molecular mechanisms of Treg induction by *E. multilocularis* remain elusive so far.

**Methodology/Principal findings:** We herein demonstrate that excretory/secretory (E/S) products of the *E. multilocularis* metacestode promote the formation of Foxp3^+^ Treg from CD4^+^ T-cells *in vitro* in a TGF-β-dependent manner. We also show that host T-cells secrete elevated levels of the immunosuppressive cytokine IL-10 in response to metacestode E/S products. Within the E/S fraction of the metacestode we identified an *E. multilocularis* activin A homolog (EmACT) that displays significant similarities to mammalian Transforming Growth Factor-β (TGF-β)/activin subfamily members. EmACT obtained from heterologous expression promoted host TGF-β-driven CD4^+^ Foxp3^+^ Treg conversion *in vitro*. Furthermore, like in the case of metacestode E/S products, EmACT-treated CD4^+^ T-cells secreted higher levels of IL-10. These observations suggest a contribution of EmACT in the *in vitro* expansion of Foxp3^+^ Treg by the *E. multilocularis* metacestode. Using infection experiments we show that intraperitoneally injected metacestode tissue expands host Foxp3^+^ Treg, confirming the expansion of this cell type *in vivo* during parasite establishment.

**Conclusions/Significance:** In conclusion, we herein show that *E. multilocularis* larvae secrete a factor with clear structural and functional homologies to mammalian activin A. Like its mammalian homolog, this protein induces the secretion of IL-10 by T-cells and contributes to the expansion of TGF-β-driven Foxp3^+^ Treg, a cell type that has been reported crucial for generating a tolerogenic environment to support parasite establishment and proliferation.

**AUTHOR SUMMARY:** The metacestode larval stage of the tapeworm *E. multilocularis* grows infiltratively, like a malignant tumor, within the organs of its human host, thus causing the lethal disease alveolar echinococcosis (AE). Immunosuppression plays an important role in both survival and proliferation of the metacestode, which mainly depends on factors that are released by the parasite. These parasite-derived molecules are potential targets for developing new anti-echinococcosis drugs and/or improving the effectiveness of current therapies. Additionally, an optimized use of such factors could help minimize pathologies resulting from over-reactive immune responses, like allergies and autoimmune diseases. The authors herein demonstrate that the *E. multilocularis* metacestode releases a protein, EmACT, with significant homology to activin A, a cytokine that might support host TGF-β in its ability to induce the generation of immunosuppressive regulatory T-cells (Treg) in mammals. Like its mammalian counterpart, EmACT was associated with the expansion of TGF-β-induced Treg and stimulated the release of elevated amounts of immunosuppressive IL-10 by CD4+ T-cells. The authors also demonstrate that Treg are locally expanded by the metacestode during an infection of mice. These data confirm an important role of Treg for parasite establishment and growth during AE and suggest a potential role of EmACT in the expansion of these immunosuppressive cells around the parasite.

## INTRODUCTION

The metacestode larval stage of the fox-tapeworm *Echinococcus multilocularis* is the causative agent of alveolar echinococcosis (AE), one of the most dangerous zoonoses world-wide [1,2]. Intermediate hosts (rodents and, occasionally, humans) usually get infected by oral ingestion of infectious eggs that contain the oncosphere larval stage. Upon hatching in the small intestine and penetration of the intestinal wall, the oncosphere gains access to the host organs and, almost exclusively within the liver, develops into the cyst-like metacestode, following a process of stem cell-driven metamorphosis [3,4]. The multi-vesicular *E. multilocularis* metacestode tissue subsequently grows infiltratively, like a malignant tumor, into the surrounding host tissue, eventually leading to organ failure and host death [2]. In later stages of the disease, metastases can be formed in secondary organs, which is probably due to the distribution of parasite stem cells via bloodstream and the lymphatic system [3]. In mice, the initial establishment phase of the parasite (the oncosphere-metacestode transition) is typically accompanied by a potentially parasitocidal, Th1-dominated immune response which, in permissive hosts, is skewed towards a permissive Th2-dominated immune response during the chronic phase of the disease [5]. Current treatment options against AE are very limited and include surgery, which can only be applied in few cases, and/or chemotherapy with benzimidazoles [2]. However, due to significant adverse side effects, only parasitostatic doses of these compounds can be applied and, consequently, the drugs often have to be administered lifelong [2]. These limitations in current AE therapy underscore an urgent need for the development of novel anti-parasitic measures.

During asexual multiplication, the *E. multilocularis* metacestode tissue persists for prolonged periods of time in close contact to immune effector cells without being expelled by the host immune response [5]. Immune suppressive mechanisms, provoked by parasite surface structures and/or excretory/secretory (E/S) products, have thus been proposed to support long-term persistence of the parasite within the host [5–8]. Accordingly, PBMCs of patients with active AE and host cells in the vicinity of parasite liver lesions in mice have been demonstrated to produce elevated levels of the immunosuppressive cytokines TGF-β and IL-10 and are believed to play important roles in the pathophysiology of AE [9–11]. Furthermore, immune effector cells from *E. multilocularis*-infected hosts typically display impaired immune reactivity [9,12–16] whereas those from hosts with degenerating parasite tend to recover their immune potential [17]. Moreover, host immune-stimulation during an infection can lead to considerably reduced disease progression [18,19]. Although the molecular and immunological basis for the immune suppression in AE is largely elusive so far, parasitic helminths as a whole have repeatedly been reported to exploit the host immune system’s own self-regulatory signaling pathways for successful establishment of an infection and long-term persistence within the host [20].

Of particular importance for regulation in mammalian immune responses are signals delivered by TGF-β superfamily members. On the basis of sequence similarities, two cytokine sub-families can be distinguished within this superfamily: the TGF-β/activin sub-family and the Bone Morphogenetic Protein (BMP) sub-family [20]. The former subfamily has gathered considerable interest concerning mechanisms of immune homeostasis maintenance [21,22]. Produced as large pro-forms consisting of an N-terminal signal peptide, followed by a pro-peptide separated by a furin recognition motif from the C-terminal active peptide (∼15 kDa), TGF-β/activin ligands are secreted as dimers of their active peptide, following cleavage of the signal sequence and pro-peptides [23]. Two particularly relevant peptides within the TGF-β/activin subfamily, TGF-β1 and activin A (i.e. inhibin beta A homodimers), have drawn considerable attention in the search for mechanisms that lead to an impairment of immune effector cell functions and, ultimately, to an expansion of tolerogenic cells [21,22]. Both cytokines were reported to impair the function of dendritic cells (DC), NK cells, macrophages, and T-cells, and stimulate the expansion of regulatory DC and T-cells [21,22].

During echinococcosis, the impaired host immune response is paralleled by an increased expression of TGF-β signaling components in periparasitic host cells and tissues [10,11,16,24–27] with the expansion of tolerogenic CD4^+^ CD25^+^ Foxp3^+^ Treg cells [6,15,28–30]. *Echinococcus* antigens can stimulate the expression of CD25 by CD4^+^ T helper cells from AE infected patients, contributing to the differentiation into Treg [31]. Using a murine system of intraperitoneal AE (secondary echinococcosis), Mejri *et al.* [15] reported increased percentages of CD4^+^CD25^+^ T-cells in the peritoneum of *E. multilocularis* infected mice at an advanced (chronic) stage of the disease, when compared to non-infected mice. This group also found Foxp3 gene expression to be elevated in these cells and a higher frequency of CD4^+^CD25^+^Foxp3^+^ Treg cells in the peritoneum and the spleen of *E. multilocularis*-infected mice [15,28]. Subsequent studies then convincingly revealed Foxp3+ Treg as key players in the immunoregulatory processes that facilitate the establishment and persistence of *E. multilocularis* metacestode in their mammalian hosts [28–30]. Consistent with such observations, our previous report of the ability of *E. multilocularis* metacestode E/S products to expand host Treg *in vitro* [6], ultimately suggested that Treg expansion during AE, as increasingly reported in the literature, could go beyond a simple homeostatic balancing mechanism.

In the present study, we have specifically followed up on these observations to further investigate the ability of the *E. multilocularis* metacestode to increase host Tregs. We report on a parasite TGF-β superfamily ligand, EmACT (*<u>E</u>. <u>m</u>ultilocularis* <u>Act</u>ivin), which is released by the metacestode, and which promotes the ability of host TGF-beta to induce Treg conversion and the production of IL-10 by host T-cells. Our data support a role of parasite-derived factors in the impairment of host immune response during AE and identify EmACT as a potent driver of host immune suppression by the *E. multilocularis* metacestode.

## METHODS

### Animals and Ethics statement

Wild type C57Bl/6 mice and Mongolian jirds were purchased from Charles River and housed at the local animal facilities of the Institute of Hygiene and Microbiology and the Institute for Virology and Immunobiology of the University of Würzburg (Germany) at least 1-2 weeks before experimentation. OT-II mice (TCR transgenic mice where CD4^+^ T cells are specific for I-A^b^ presentation of OVA_323–339_ peptide) were kindly provided by Francis Carbone, Melbourne, Australia. OT-II mice were crossed with C57Bl/6 Rag-1^−/−^ mice (devoid of mature B and T-cells), a generous gift from Thomas Winkler, Erlangen, Germany. All animal handling, care and subsequent experimentation were performed in compliance with European and German regulations on the protection of animals (*Tierschutzgesetz*). Ethical approval of the study was obtained from the local ethics committee of the government of Lower Franconia (Regierung von Unterfranken, 55.2-2531.01-31/10 and 55.2-2531.01-26/13).

### *In vitro* maintenance of *E. multilocularis* metacestode and collection of E/S products

*E. multilocularis* metacestodes were isolated, separated from host contaminants and axenically cultivated as previously described [6]. For the collection of E/S products, axenically maintained metacestode vesicles were washed thrice in 1 × PBS and resuspended in collection medium DMEM10redox i.e. Dulbeccós Modified Eaglés Medium, 4.5 g glucose/L (DMEM ^+^ GlutamaxTM, GIBCO) supplemented with 10% Fetal Bovine Serum Superior (Biochrom AG), 100µg/ml penicillin/streptomycin (PenStrep solution, Biochrom AG), 20 µg/ml Levofloxacin (Tavanic, Sanofi-Aventis, Deutschland GmbH), 143 µM β-mercapthoethanol (Sigma-Aldrich, cat. M6250), 10 µM Bathocuproine disulfonic acid (Sigma, cat. B-1125) and 100 µM L-Cysteine (Sigma, cat. C-1276) under axenic conditions (i.e. sealed in Nitrogen filled Ziploc freezer bag and placed in a 5%CO2 incubator at 37°C). After 48 hours of culture, the supernatants containing the metacestode E/S products were collected and filtered through a 0.2 μm sieve (Filtropur S filter, SARSTEDT). The total amount of E/S product proteins was determined using the bicinchoninic acid assay (Pierce BCA Protein Assay Kit, ThermoScientific, prod # 23228) and the E/S products stored at −80°C until use.

### Injection of *E. multilocularis* metacestodes and *in vivo* follow-up

#### Preparation of parasite material and injections

For *in vivo* assays, metacestode vesicles were obtained from infected Mongolian jirds (*Meriones unguiculatus*). The recovered parasite homogenates were washed thrice in PBS (1X) then transferred to DMEM10redox for axenic maintenance with medium change twice per week for up to 10 days. The complete removal of host contaminants was assessed by PCR as previously described [6]. The host-free parasite homogenates were then washed in PBS (1X), separated in aliquots of 5000 acephalic cysts resuspended in a total volume of 500 µl PBS (1X) to be used for intraperitoneal injections. An equal volume (500 µl) of the carrier solution PBS (1X) was used in parallel for mock injections. Parasite preparations from 5 unrelated iird infections were used to include any eventual parasite-related variation in the analysis.

#### Peritoneal lavage and cell collection

At defined points within a time frame of 42 days post intraperitoneal injection, mice were sacrificed by CO_2_ asphyxiation. Ice cold PBS (1x) with 10% heat-inactivated filtered FBS Superior (Biochrom AG) was then used to wash the peritonea and retrieve the peritoneal exudate cells. In each of the five experimental replicates (injections performed using five different isolates), the peritoneal cells were harvested from naïve (a pool of 3 mice) or infected (1 mouse) mice at each time point. The suspensions were filtered through a 70µm nylon cell strainer (BD Biosciences). Red blood cells in the filtrates were lysed with 1.4% NH4Cl for 5 minutes at 37°C. The filtrates were then washed in R10 medium and the total numbers of recovered cells determined using the trypan blue (Biochrom) exclusion test on a bright-line Neubauer counting chamber prior to analysis. At the end of the infection follow up (42 days), the parasite tissues were thoroughly harvested from the peritonea of each infected mice and the masses were determined.

#### Flow cytometry

2 × 10^5^ peritoneal exudates cells were stained with fluorochrome-conjugated antibodies (anti-mouse) directed against the T-cell subset surface marker CD4 (Biotin, Miltenyi Biotec), the alpha chain of the IL-2 receptor CD25 (PE, eBioscience) present on activated T-cells and the intracellular master transcription factor of Treg, Foxp3 (APC, Miltenyi Biotec). Biotinylated CD4 antibodies were detected by incubation with either FITC- or Pe-Cy5-conjugated streptavidin (BD Biosciences). As isotype control of activated/regulatory T-cells, Rat IgG1 K Isotype (PE, eBioscience) was used. The staining procedure was executed as per the manufacturer instructions (Treg Detection Kit, Miltenyi Biotec). The cells were resuspended in FACS buffer (1x PBS supplemented with 3% heat-inactivated and filtered FCS and 0.1%NaN3) and acquired on a cytometer (FACSCalibur, Beckton Dickinson) equipped with CellQuest software. Results were further analyzed on FlowJo software (Tree Star, USA).

### *In vitro* Treg suppression assay

From the spleen of a healthy mouse, and peritoneal exudates cells of mice 7 days post infection (20 mice pooled), CD4^+^ cells were separately isolated using mouse CD4^+^ T-cell negative selection protocol (EasySep™ Mouse CD4^+^ T-cell Enrichment Kit, Stem Cell Technologies) to a purity of > 90% according to the manufactureŕs instructions. Purified CD4^+^ T-cells were stained with CD4 antibody (Biotin, Miltenyi Biotec) and CD25 antibody (PE, eBioscience) followed by incubation with Pe-Cy5-conjugated streptavidin (BD Biosciences). CD4^+^CD25^−^ and CD4^+^CD25^+^ cells were then sorted on a MoFlo high-speed sorter (Cytomation). Sorted CD4^+^CD25^−^ splenic cells (responders) were labeled with 2µM CFSE (CFDA SE, Molecular Probes/Invitrogen) at 37°C for 10 min and washed twice in R10 medium before use. For *in vitro* proliferation of the isolated responder cells, total splenocytes from a healthy mouse, were labeled with a cocktail of non-APC binding antibodies namely the murine T-cell lineage directed anti-Thy-1.2 antibody (BD Pharmingen) and the T-cell subsets recognizing CD4 antibody (eBioscience) and CD8 antibody (eBioscience) on Ice for 30 minutes. The cells were then washed in 1x PBS supplemented with 3% heat-inactivated and filtered Fetal Calf Serum (FCS, PAA Laboratories) prior to antibody mediated complement lysis (Rabbit complement, Cedarlane) at 1/10 dilution for 45 minutes under constant agitation at 37°C. Next, the suspension was filtered through a 70 µm nylon cell strainer (BD Biosciences) and the filtrate, representing antigen presenting cells (APC), washed and resuspended in R10 medium (RPMI-1640 from GIBCO BRL supplemented with 100U/ml Penicillin (Sigma), 100µg/ml Streptomycin (Sigma), 2mM L-glutamin (Sigma), 50 µM β-mercaptoethanol (Sigma) and 10% heat-inactivated and filtered (0.22 µm, Millipore) fetal calf serum (FCS, PAA Laboratories). APC were then irradiated on an X-ray unit (Faxitron, CellRad) with 20 Grays and counted using the trypan blue (Biochrom) exclusion test on a bright-line Neubauer counting chamber. A total of 2 × 10^5^ irradiated APCs was then cultured in CD3 antibody (1ug/ml, eBioscience) pre-coated 96-well round-bottom plates with 2 × 10^4^ CFSE-labelled responders (Splenic CD4^+^CD25^−^ cells) and 1-2 × 10^4^ CD4^+^CD25^+^ cells for 5 days. The cells were then harvested, resuspended in FACS buffer (1x PBS supplemented with 3% heat-inactivated and filtered FCS and 0.1% NaN3) prior to acquisition on a cytometer (FACSCalibur, Beckton Dickinson) equipped with CellQuest software. Results were further analyzed on FlowJo software (Tree Star, USA).

### Identification, cloning and analysis of the Em*act* cDNA and gene

The full length sequence of the *Schistosoma mansoni* TGF-β/activin subfamily member (SmInAct, DQ863513) and the human inhibin beta A chain (HsINHßA, P08476) were used to search the *E. multilocularis* genome on Wormbase (https://parasite.wormbase.org/Echinococcus_multilocularis_prjeb122/Info/Index/) using t*blast*n and *blastp* algorithms. A predicted gene with a truncated 5’end (EmuJ_000178100) could be retrieved from the available *E. multilocularis* genome sequence [32]. The full-length coding sequence of the corresponding cDNA was identified by screening of a complementary DNA library [33] and termed *Emact*. Briefly, a consensus sequence between the *E. multilocularis* genome scaffold 6 and Sm*Inact* was used as template for primer design. The following primers were designed and used for retrieval of the 5’ (*Emact*_5’: 5’-ACA GTA GTT GGG TTC-3’) and 3’ (*Emact*_3’: 5’-GAA CCC AAC TAC TGT-3’) ends of Em*act*. These primers were used in pairs with primers specifically recognizing the carrier vector part of the cDNA library, pJG4-5 [34]. Once recovered, the 5’and 3’ends of the parasite putative *act* reading frame, were used to design primers for the full length amplification of the *Emact* coding sequence, namely *Emact*_Dw (5′-ATG ACC ATT ACT ACC CCC ATG AAG-3′) and *Emact*_Up (5’-ACT ACA ACC GCA CTC TAG GAC AAT G-3’). Metacestode RNA was isolated using Trizol reagent (Invitrogen) and 1µg of total RNA was reverse transcribed with Omniscript RT kit (Qiagen) according to the manufacturerś instructions. The generated cDNA was used as template for amplification of the *emact* full transcript using the primer pair *Emact*_Dw x *Emact*_Up by high fidelity polymerase chain reaction (Phusion, NEB). Resulting amplicons were sub-cloned into the pDrive cloning vector (QIAGEN) and five clones were picked and sequenced in both directions identically revealing the full coding sequence of *emact* (EmuJ_000178100). Sequence similarities between the deduced amino acid sequence of *Emact* and other members of the TGF-β superfamily were determined through multiple sequence alignments using BIOEDIT (http://www.mbio.ncsu.edu/BioEdit/bioedit.html), and a neighbor-joining tree was generated from alignments using MEGA [34].

### EmACT Antibody production

For the production of polyclonal antibodies, EmACT was expressed in the bacterial pBAD/TOPO ThioFusion Expression Kit (Invitrogen). To maximize the recognition of EmACT after processing by the generated antibodies, full *Emact* (without stop codon) amplified using the primer pair *Emact*_Dw / *Emact*_Up coding for the preproprotein EmACT was chosen for immunization and subcloned in pBAD/TOPO ThioFusion expression vector (Invitrogen). The thioredoxin-fusion protein (Thio-EmACT) with histidine tag, expressed in *E.coli* Top10 cells by adding arabinose (2 g/L; 4 hours), was purified on nickel-nitrilotriacetic acid resin (Invitrogen) according to the manufactureŕs protocol. The purified Thio-EmACT was then diafiltered on Centrifugal Filter Units (Millipore) against sterile PBS (1X) before quantification using the BCA assay (ThermoScientific). NMRI mice were injected subcutaneously at two different locations with 100µg of the recombinant Thio-EmACT resuspended in 100µl Freund Incomplete adjuvant (Sigma). The double injections were repeated four weeks later to boost the mice anti-EmACT response. Finally, ten days after the boost, the mice were bleeded from the heart and the serum collected and stored at −20°C until use. In parallel, naïve mice were also bleeded and the serum collected and stored as normal mouse serum.

### Recombinant expression of EmACT in Human Embryonic Kidney cell line

Em*act* (without signal peptide) was subcloned in the eukaryotic pSecTag2 expression system (Invitrogen) to generate the pSegTag2-Em*act* vector construct as per the manufacturer instructions. Human embryonic kidney cell-line 293T (HEK 293T) were transfected with the expression vector construct pSegTag2-Em*act* or the empty pSecTag2 vector (Invitrogen) as control. Transfections were performed using linear polyethyleneimine (25 kDa, Sigma) according to the manufactureŕs instructions. All transfections were performed in petri dish (92 × 16 mm [Ø × height], SARSTEDT). HEK cells were seeded 16 hours prior to transfection (3 × 10^6^ cells / dish). 24 hours post-transfection, the supernatants were replaced with fresh DMEM10 medium (i.e. Dulbeccós Modified Eaglés Medium, 4.5g glucose/L (DMEM ^+^ Glutamax, GIBCO) supplemented with 10% Fetal Bovine Serum Superior (Biochrom AG), 100µg/ml Penicillin/streptomycin (PenStrep solution, Biochrom AG) and 20µg/ml Levofloxacin (Tavanic, Sanofi-Aventis, Deutschland GmbH).). The supernatants of transfected HEK cells were then collected after 24 hours of incubation, filtered over a bottle top filter (Filtropur BT50, SARSTEDT), normalized for the total protein content (BCA Protein Assay Kit, ThermoScientific) and stored as aliquots at −80°C until use.

### Immunodetection

To detect EmACT in supernatants of parasite cultures (natural) or HEK 293 T cell cultures (recombinant), 1ml of metacestode vesicle E/S products (MVE/S) or pSecTag2-Em*act*-transfected HEK cell supernatant was resuspended in 9 volumes of 100% ice-cold ethanol. The mixtures were kept at −80° C for at least 2 hours, and then centrifuged for 30 min at 14,000 rpm in a refrigerated centrifuge. The supernatants were discarded and the pellets were dried thoroughly at 50°C and resuspended in 50 µl of 2 × STOPP mix (2 ml 0.5M Tris-HCl pH 6.8, 1.6ml glycerol, 1.6ml 20% SDS, 1.4 ml H2O, 0.4 ml 0.05% (w/v) bromophenol blue, 7 µl β-mercaptoethanol per 100 µl) and boiled for 10 min at 100°C. 10µl of each protein sample were separated by SDS-PAGE and transferred to a nitrocellulose membrane for immunodetection with anti-EmACT antiserum or with mouse pre-immune serum.

### Generation of murine bone marrow-derived dendritic cells (BMDC)

Dendritic cells were obtained by GMCSF-driven differentiation of mice bone marrow precursor cells as previously described [35]. Briefly, C57Bl/6 mice (Charles River/Wiga, Sulzfeld, Germany) aged 6-14 weeks and bred within the animal facility of the Institute of Virology and Immunobiology, University of Würzburg, under specific pathogen-free conditions were sacrificed using CO_2_ asphyxiation. Femurs and tibiae were removed and purified from the surrounding muscle tissue. Thereafter the marrow was flushed with PBS (1X), resuspended by gently pipetting and washed once in medium. The medium used here was R10 medium composed of RPMI-1640 (GIBCO BRL) supplemented with 100 U/ml Penicillin (Sigma), 100 µg/ml Streptomycin (Sigma), 2 mM L-glutamine (Sigma), 50 µM β-mercaptoethanol (Sigma) and 10% heat-inactivated and filtered (0.22 µm, Millipore) fetal calf serum (FCS, PAA Laboratories) as previously defined. Once resuspended, the bone marrow precursor cells were counted using the trypan blue (Biochrom) exclusion test on a bright-line Neubauer counting chamber. 2 × 10^6^ precursor cells were cultured in R10 medium supplemented with 10% GMCSF-containing supernatant as previously defined [35]. At day 8, non-adherent cells representing at a high frequency newly differentiated dendritic cells (70-89% CD11c^+^) were harvested, washed once in R10 medium prior to subsequent assays.

### Isolation of splenocytes and lymph node cells

Single cell suspensions were obtained from the spleen and lymph nodes of C57Bl/6 mice by mechanically squeezing the tissues with glass slides in cold PBS and filtered through a 70 µm nylon cell strainer. Red blood cells in the spleen filtrate were lysed with 1,4% NH_4_Cl for 5 min at 37°C, and the splenocytes were washed in R10 medium, that is RPMI 1640 (GIBCO BRL) supplemented with penicillin (100 U/ml, Sigma, Deisenhofen, Germany), streptomycin (100 μg/ml, Sigma), L-glutamin (2 mM, Sigma), 2-mercaptoethanol (50 μM, Sigma and 10% heat-inactivated fetal calf serum (FCS, PAA Laboratories, Parsching, Austria). Cell counts were subsequently determined for splenocytes and lymph node cells using the trypan blue (No.26323, Biochrom, Berlin, Germany) exclusion test on a bright-lined Neubauer counting chamber.

### Treg conversion assays

CD4+ T-cells were isolated from murine splenocytes and lymph node cells using a T-cell negative selection kit (Easy Sep mouse T-cell enrichment kit, Stem Cell Technologies) to a purity >90% as per the manufactureŕs instructions. CD4^+^ T-cells were further enriched for CD25^−^ cells using Miltenyi Biotećs LD columns with a suitable MACS separator achieving > 90% purity as per the manufactureŕs instructions. Murine BMDCs at day 8 of cultivation were incubated with 3-fold higher numbers of CD4^+^ CD25^−^ T-cells (OT-II or OT-II.RAG-1^-/-^) and 200 ng/ml OVA peptide (323-339, grade V, Sigma) supplemented or not with our different stimuli. In some assays, the cultures were supplemented with 20 μg/ml of a pan-vertebrate anti–TGF-β blocking antibody 1D11 (R&D Systems), alongside stimuli addition. In others, isolated naïve T-cells were pre-incubated for 30 minutes with 5 μM of an inhibitor of TGF-β superfamily type I activin receptor-like kinase (ALK) receptors ALK4, ALK5, and ALK7 (SB431542 [36]) before cultivation with BMDC.

Alternatively, CD4^+^ T-cells were purified from wild type C57Bl/6 mice and subsequently activated on plate-bound CD3 (1 µg/ml) and CD28 (0.5 µg/ml) antibodies in the absence or presence of our different stimuli. Recombinant human TGF-β1 (R&D Systems) was used as positive control. After 5 days of incubation, the cells were harvested and stained using the Treg detection kit (Miltenyi Biotec), resuspended in FACS buffer (1x PBS supplemented with 3% heat-inactivated and filtered FCS and 0.1%NaN3) prior to acquisition on a cytometer (FACSCalibur, Beckton Dickinson) equipped with CellQuest software. Results were further analyzed on FlowJo software (Tree Star, USA).

### CD4^+^ T-cells stimulation assay

CD4^+^ T-cells were isolated from splenocytes and lymph node cells of wild type C57Bl/6 mice (6-8weeks old) using a T-cell negative selection kit (Easy Sep CD4^+^ T-cell enrichment kit, Stem Cell Technologies) to > 90% purity according to the manufactureŕs instructions. The CD25^−^ fraction was further enriched using Miltenyi Biotec LD columns with a suitable MACS separator achieving > 90% purity for CD4^+^CD25^−^ T-cells. Next, 2 × 10^5^ CD4^+^CD25^−^ T-cells were seeded in a 24-well tissue culture plate (Flat bottom, SARSTEDT) that had been coated with CD3 antibody (0.1µg/ml, eBioscience) overnight at 4°C in R10 medium. The cells suspension was supplemented with 5µg/ml of CD28 antibody (eBioscience) and different stimuli were subsequently added. After 72h, T-cells supernatants were probed for IL-10 release by ELISA as per the manufactureŕs instructions (BD OptEIA - Mouse IL-10 ELISA Set - BD Biosciences with a detection limit of 19pg/ml).

### Statistical Analyses

All results were expressed as mean ± standard deviation (SD). Differences observed between groups were evaluated using the Wilcoxon/Mann-Whitney U test, a nonparametric test that does not assume normality of the measurements (it compares medians instead of means). Values of p<0.05 were considered statistically significant. Statistical analyses were performed with a statistical software analyzing package (GraphPad Software).

### Accession numbers

The complete *Emact* cDNA sequence reported in this paper was deposited in the GenBank database under the accession number HF912278. All GenBank accession numbers of genes and sequences used in this study are listed in S1 Tab.

## RESULTS

### *E. multilocularis* metacestode tissue drives Foxp3^+^ Treg expansion in experimentally infected mice

In previous reports, it has been shown that Treg are expanded during chronic secondary AE 6-12 weeks post infection [15,28]. It has not been elucidated, however, whether this expansion resulted from an inherent host protective mechanism in the course of a chronic infection in order to minimize tissue damage, or whether Treg expansion was actively driven by the parasite. Since chronic AE, other than early AE, is associated with severe depletion of T-cells after 6 weeks of infection [37], we investigated the dynamics of host CD4^+^ T-cell responses during experimental secondary AE for up to 7 weeks (42 days) post infection; i.e. up to the chronic phase of the disease. To this end, we injected mice intraperitoneally with 5000 *E. multilocularis* acephalic cysts (metacestode), axenically sub-cultured for up to 10 days to remove host cells (as confirmed by species-specific PCR, see Fig 1A), and analyzed the peritoneal exudate cells over a 7-weeks period (42 days).

**Fig 1:**
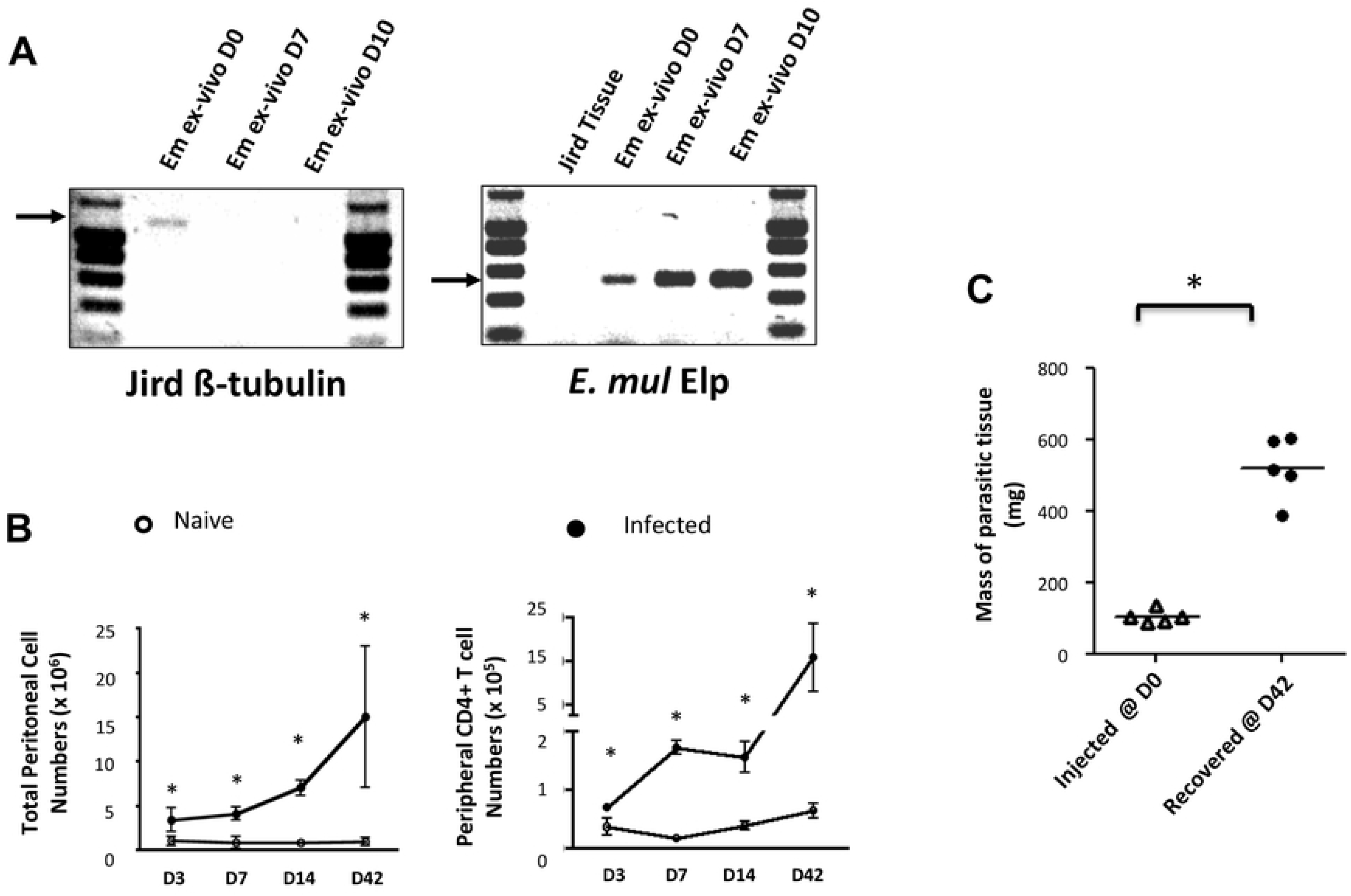
Injected *E. multilocularis* metacestode tissue proliferates in the face of accumulating CD4^+^ T-cells. (**A**) Parasite cysts were harvested from infected jirds and kept under axenic conditions for 10 days. The presence of host contaminants was assessed by organism-specific PCR. Chromosomal DNA was isolated, the host (Jird)-specific ß-tubulin or the parasite-specific *elp* genes were separately amplified. Jird tissue was used as a negative control for the parasite-specific gene *elp*. (**B**) Peritoneal exudate cells were collected, counted and analyzed by flow cytometry for CD4 expression. Parasite-driven accumulation of total (**B** up) or CD4^+^ T-cells (**B** down) is shown for D3-42 post injection. (**C**) Masses of parasitic tissue injected and recovered after 42 days. Horizontal bars stand for mean levels. Data represent means +- SD from groups of five mice for each time point assayed individually (Infected). Naive mice were clustered in sub-groups of 3 mice pooled as one per assay (15 mice for each time point).*, *p* < 0.05.

We observed a significant increase of peritoneal exudate total and CD4^+^ T-cells over time in *E. multilocularis* infected mice as compared to mock (PBS)-injected controls (Fig 1B). Interestingly, despite the anti-AE role of host cellular immunity in general and CD4^+^ T-cell mediated effector functions in particular [38], the CD4^+^ T-cell expansion in infected mice was paralleled by an increase in parasite mass during the study period (Figure 1C).

We then examined the subsets of CD4^+^ T-cells expanded in infected mice by expression levels of CD25 and Foxp3. A separation into CD25^+^Foxp3^−^ CD4^+^ activated effector T-cells (Teffs) and CD25^+^Foxp3^+^ CD4^+^ T-cells as Tregs was applied (Fig 2A). Although we noted a general increase of Foxp3^+^ Treg numbers in infected mice when compared to naïve mice throughout the study period (Fig 2B), a transient but significant increase of the proportion of Tregs was uniquely detectable at 7 days post-infection within the peritoneal exudates of mice (Fig 2C). The Tregs induced by the parasite 7 days post intraperitoneal inoculation were able to repress proliferative responses of CFSE-labeled conventional CD4^+^CD25^−^ T-cells (Fig 3), indicating that they were functionally suppressive.

**Fig 2:**
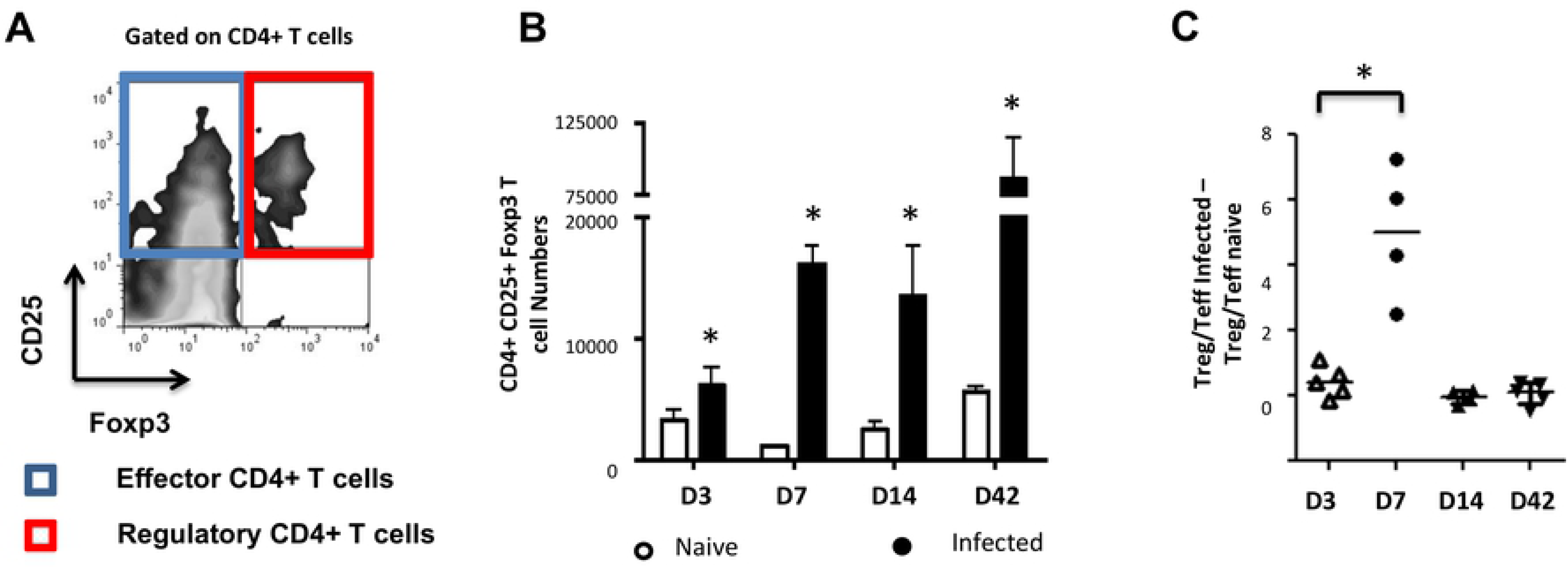
Initial CD4^+^ T-cell responses to *E. multilocularis* metacestodes are biased towards Foxp3^+^ Treg. The effects of intraperitoneal injection of *E. multilocularis* cysts in C57Bl/6 mice were followed by analysis of peritoneal exudate CD4^+^ cells harvested at days 3, 7, 14 and 42 post-injection, respectively. The cells were analyzed by flow cytometric analysis for CD25 and Foxp3 expression. (**A**) The CD25^+^ population was clustered into Teffs or Tregs with regards to Foxp3 expression. (**B**) The kinetics of Foxp3^+^ Treg numbers was monitored. A Mann Withney U test was performed separately at each time point to compare naive and infected mice. (**C**) Kinetics of Treg/Teff ratio over time as a measure of the bias of parasite-associated CD4^+^ T-cell response. Each ratio for infected mice was substracted of the corresponding naive mice ratio. Data represent means +- SD from groups of five mice for each time point assayed individually (Infected). Naive mice were clustered in sub-groups of 3 mice pooled as one per assay (15 mice for each time point).*, *p* < 0.05.

**Fig 3:**
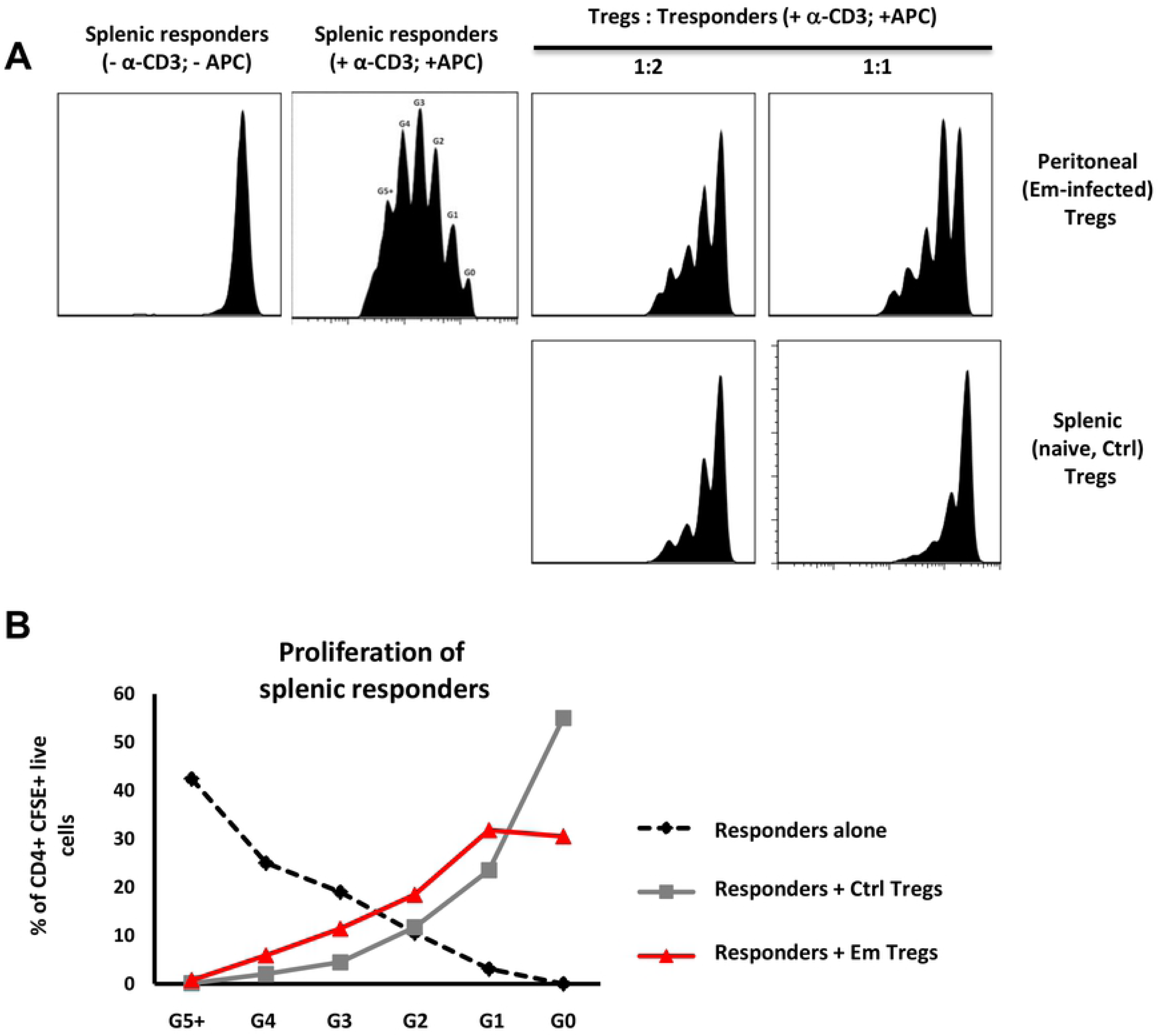
*E. multilocularis* metacestode-induced Treg are functionally suppressive *in vitro*. Peritoneal exudate cells from mice infected for 7 days with 5000 acephalic *E. multilocularis* cysts, and naive splenocytes from control mice, were prepared by CD4^+^ T-cell magnetic selection, then FACS-sorted into CD4^+^CD25^+^ and CD4^+^CD25^−^populations. Splenic naive CD4^+^CD25^−^cells (responders) were then polyclonally stimulated in the presence or absence of CD4^+^CD25^+^ cells from either control or *E. multilocularis*-infected mice. (**A**) Representative plots of CFSE-labelled responder cells proliferation with increasing amounts of CD4^+^ CD25^+^ Treg. (**B**) Proportions of labeled live CD4^+^ cells in each generation of assay conducted at 1:1 ratio, as gated by CFSE dilution. A representative experiment out of two with similar results is displayed.

Taken together these analyses showed that *E. multilocularis* metacestodes can grow in these mice and raise a CD4^+^ T-cell response but with a transient overproportional expansion of suppressive Tregs.

### E/S products of the *E. multilocularis* metacestode induce Foxp3 expression and IL-10 production by host T-cells *in vitro*

We previously demonstrated Foxp3^+^ Treg expansion *in vitro* from OT-II naïve CD4^+^ T-cells activated with OVA-loaded DC in the presence of *E multilocularis* metacestode E/S products [6], suggesting either a direct induction of Treg conversion by the parasite products or a mitogenic effect of these products on pre-existing OT-II Treg. To further examine these alternatives, we isolated naïve OT-II.RAG-1^-/-^ CD4^+^T-cells from spleens and lymph nodes of naïve animals, genetically devoid of Foxp3^+^ Tregs (Fig 4A). The cells were activated *in vitro* with OVA-loaded DCs in the presence of *E. multilocularis* metacestode E/S products (MVE/S) as previously described [6]. MVE/S failed to activate BMDC cultures beyond the baseline level obtained with medium, arguing against a potential contamination of the harvested parasite products with endotoxins [6]. Notably, Foxp3^+^ Treg frequencies were considerably enhanced in cultures supplemented with MVE/S, similar to TGF-β (Fig 4B), suggesting that *E. multilocularis* metacestode E/S products can induce *de novo* Foxp3^+^ Treg conversion from naive CD4^+^ T cells *in vitro*. We also measured the production of the immunosuppressive cytokine IL-10 in DC-T-cell co-cultures in the presence or absence of the parasite products. We noted a significantly increased production of IL-10 in cultures supplemented with *E. multilocularis* metacestode products (Fig 4C) indicating that the parasite products can both expand host Foxp3^+^ Treg, and also trigger an elevated production of IL-10 by host immune cells.

**Fig 4:**
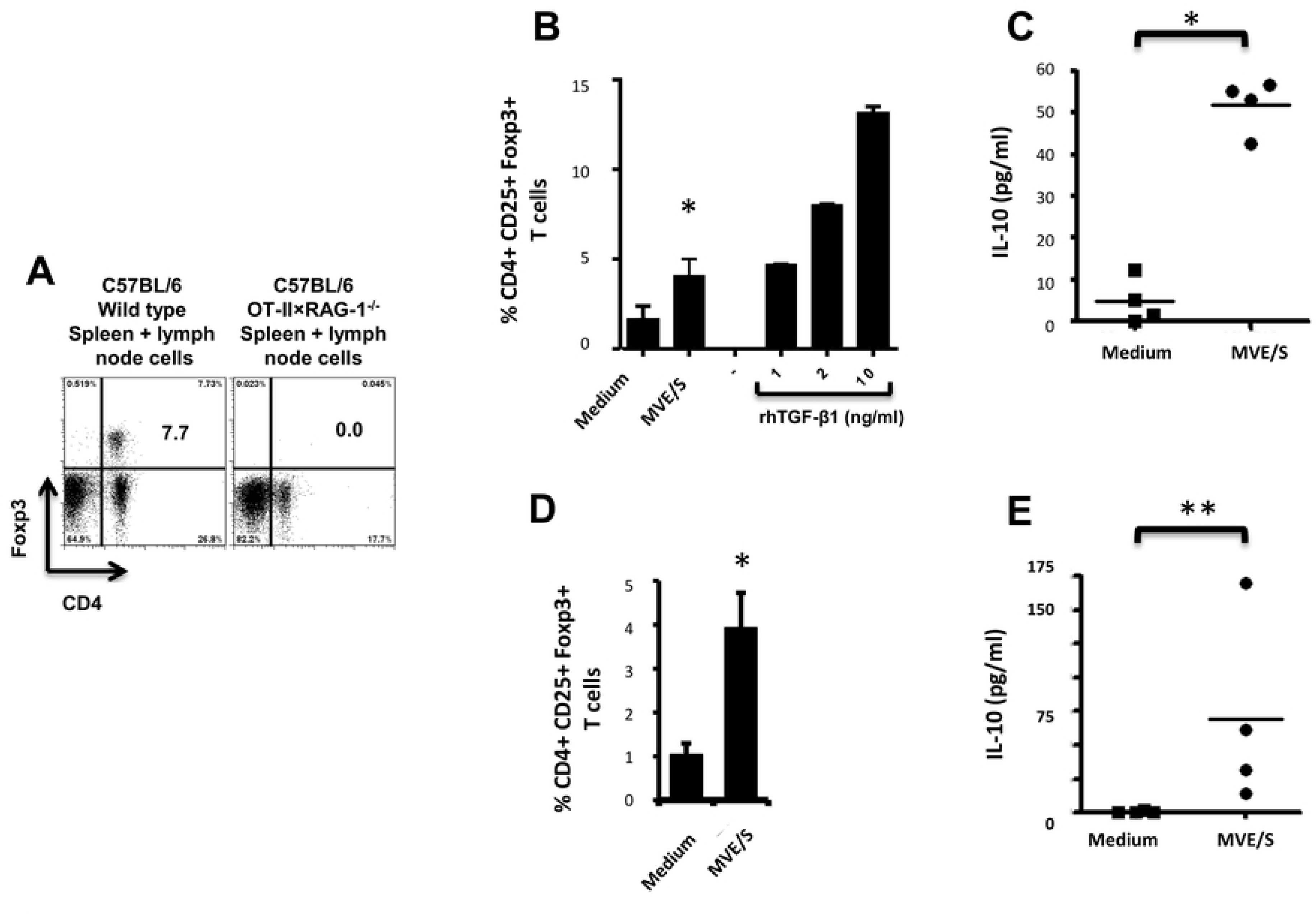
E/S products of *E. multilocularis* metacestode promote the *de novo* Foxp3^+^ Treg conversion *in vitro*. (**A**) Staining of CD4^+^ Foxp3^+^ T-cell within the bulk of spleen and lymph node cells from wild type C57Bl/6 or C57Bl/6 OT-II.RAG-1^-/-^ mice over a C57Bl/6 background. (**B**) MVE/S promote *de novo* CD4^+^CD25^+^Foxp3^+^ Treg conversion *in vitro*. Freshly generated DCs (Day 8) were co-cultured with naïve CD4^+^CD25− T-cells from OT-II.RAG-1^-/-^ mice at a DC:T-cell ratio of 1:3 in R10 medium supplemented with OVA peptide (200ng/ml). E/S-free medium (DMEM10redox) or MVE/S-containing medium was added to the cultures prior to incubation. Different doses of recombinant human TGF-β1 were used as positive controls. 5 days later, cells were harvested and stained for CD4, CD25 and Foxp3 prior to flow cytometry analysis. (**C**) Additionally, culture supernatants were collected and probed for IL-10 by ELISA. (**B, C**) Summarized in the graph are the percentages of CD25^+^ Foxp3^+^ cells within the CD4^+^ T-cell population and the production of IL-10 measured after exposure to the indicated stimuli. Data represent mean ± SD from two independent experiments with products from two different parasite isolates. **(D)** Foxp3^+^ Treg frequencies in CD4^+^ Tcells cultured for 5 days on CD3/CD28 antibody-coated plates in the presence of E/S-free medium (DMEM10redox) or MVE/S-containing medium. Bars represent the mean ± SD of results obtained with E/S products from 4 different parasite isolates. *, *p* < 0.05. **(E)** Naïve CD4^+^ CD25^−^T-cells freshly isolated from C57Bl/6 mice were stimulated at 2 × 10^5^ /ml with CD3/CD28 antibodies in the presence of parasite E/S-free cultivation medium (DMEM10redox) or MVE/S-containing medium. After 72 hours, the T-cells supernatants were collected and probed for IL-10 concentration by Elisa. Horizontal bars represent the mean from experiments conducted with E/S products from 4 different parasite isolates. *, *p* < 0.05; **, *p* < 0.005.

Next, to investigate the role of the DC population in T-cell modulation by *E. multilocularis* products, naïve CD4^+^ T-cells from spleens of C57Bl/6 mice activated with plate-bound anti-CD3 and anti-CD28 antibodies (instead of DC-based activation) in the presence of *E. multilocularis* metacestode E/S products were both analyzed for Foxp3 expression and IL-10 production (see material and methods for experimental set-ups). We observed an increased rate of Foxp3^+^ Treg (Fig 4D), and a significantly elevated production of IL-10 (Fig 4E) in host T-cell cultures indicating that *E multilocularis* metacestode products can induce Foxp3^+^ Treg conversion and trigger IL-10 release by naïve T-cells in a DC-independent manner.

### Host TGF-β and TGF-β signaling are essential for Treg conversion driven by metacestode E/S products

It has previously been shown that the conversion of CD4^+^ T-cells to Treg requires TGF-β [39] and we cannot exclude that the complex, serum-containing media required for parasite cultivation do contain this cytokine to a certain amount. To analyze whether metacestode E/S products require host TGF-β activity to promote Treg conversion, anti-TGF-β neutralizing antibodies were used. The performed assay showed a clear inhibition of Foxp3^+^ Treg conversion by metacestode E/S products when TGF-β was neutralized (Fig 5). To additionally confirm an important role of TGF-β in the ability of metacestode products to expand host Treg, the TGF-β signaling inhibitor SB431542 [36] was used. Again, we observed a drastic reduction of the rate of Foxp3^+^ Treg induced by metacestode E/S products (Fig 5). Taken together, we conclude from these studies that metacestode E/S products can induce the conversion of naive CD4^+^ T-cells into Foxp3^+^ Treg *in vitro* and that this activity depended on the presence of host TGF-β and functional TGF-β signaling in host cells.

**Fig 5:**
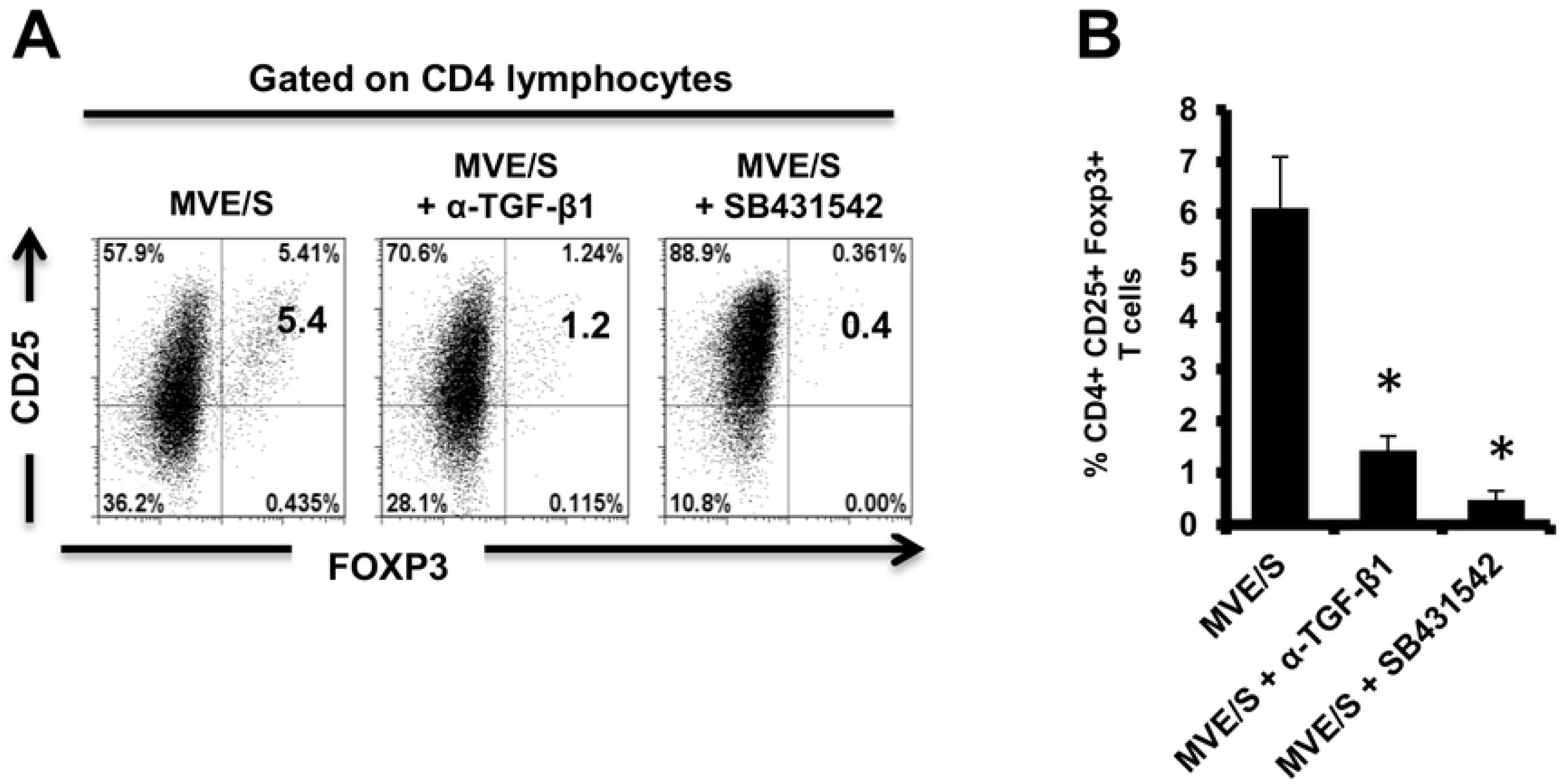
Blocking TGF-β signalling or host TGF-β alone abrogates *E. multilocularis*-driven Treg conversion *in vitro*. (**A**) Representative plots of CD25 versus Foxp3 expression, gated on CD4^+^ T-cells, from OT-II naïve CD4^+^ T-cells cultivated with freshly generated DC (Day 8) at a DC:T-cell ratio of 1:3 in R10 medium supplemented with OVA peptide (200ng/ml) in the presence of MVE/S-containing medium alone (supplemented with DMSO in one out of two experiments), combination of MVE/S-containing medium with TGF-β antibody or combination of MVE/S-containing medium with SB431542 (resuspended in DMSO). Flow cytometry was performed 5 days later. (**B**) Mean percentages of Foxp3^+^ Treg within the CD4^+^ T-cell population of DC/T-cell cultures supplemented with the indicated stimuli. Bars represent mean ± SD from two independent experiments with cells from individual mice and products from two different parasite isolates. *, *p* < 0.05.

### *E. multilocularis* expresses an activin A – like cytokine

By literature search for molecules that could exert activities as observed above for metacestode E/S products, we found striking similarities to the mammalian TGF-β-like cytokine activin A. Like metacestode E/S products, activin A can induce Treg conversion *in vitro*, which depends on host TGF-β and functional TGF-β signaling [40,41], and it can induce the secretion of IL-10 by CD4^+^ T-cells [41]. Interestingly, the expression of activin-like cytokines, SmInAct [42] and FhTLM [43], have previously been reported for the related flatworm parasites *Schistosoma mansoni* and *Fasciola hepatica*, respectively. Both of these molecules influence parasite development and immunoregulatory functions have also been demonstrated for the latter [44]. We therefore hypothesized that *E. multilocularis* might also express an activin A-like molecule and performed extensive BLASTP analyses on the published genome sequence [32], using mammalian inhibin beta A (activin A monomer) and SmInAct as queries. These analyses revealed the presence of one single-copy gene, EmuJ_000178100, encoding a protein with significant homologies to both query sequences. Since further genome mining did not yield indications for the presence of additional inhibin/activin-encoding genes, implying that the cytokine encoded by EmuJ_000178100 can only form homo-but not heterodimers, the respective gene was designated *Emact* (for *<u>E</u>. <u>m</u>ultilocularis* <u>act</u>ivin).

The full length cDNA of *Emact* was cloned and sequenced and comprised 1536 bp that encoded a 507 aa protein, EmACT, with a hydrophobic region at the N-terminus, indicating the presence of an export-directing signal peptide (Fig 6). Structurally, EmACT displayed several conserved features of the TGF-β cytokine superfamily such as a C-terminal, cysteine-rich active domain, separated by a tetrabasic RTRR cleavage motif from a large N-terminal and less well conserved pre-protein sequence. Within the C-terminal active domain of all TGF-β superfamily members (activins and BMPs) are seven invariant cysteine residues, six of which form a rigid, heat stable “cysteine knot” [23]. Accordingly, the C-terminal domain (130 aa) of EmACT contained all these invariant cysteines (Fig 6, Fig 7), as well as two additional cysteines (Fig 6, Fig 7) that are characteristic of TGF-β/activin subfamily members, but that are not present in BMP subfamily members (Fig 7).

**Fig 6:**
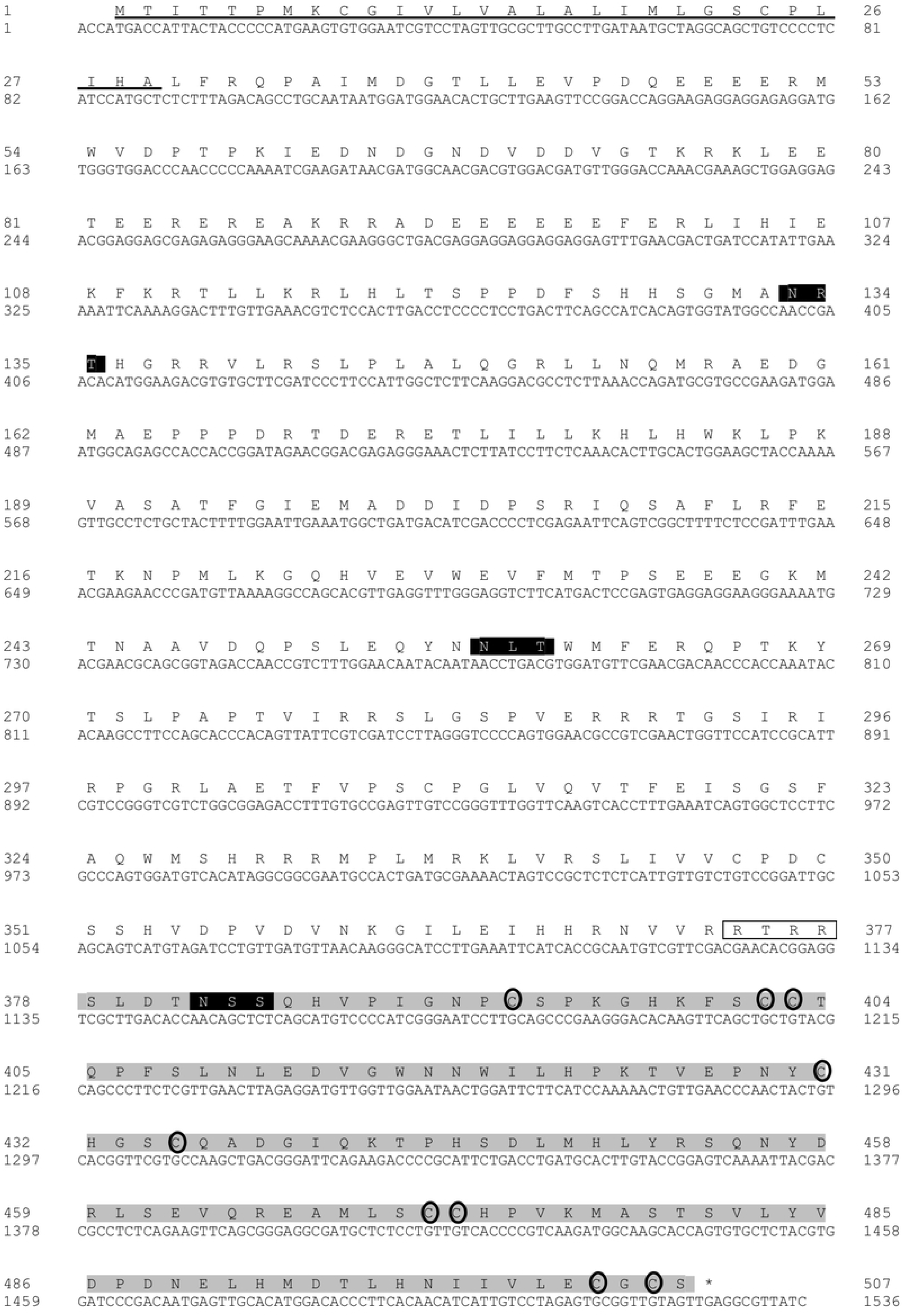
Nucleotide and deduced amino acid sequence of the *Echinococcus multilocularis act* cDNA. The 5’signal sequence is underlined. The potential N-glycosylation sites (NRT, NLT and NSS) are shown in solid boxes. The paired dibasic cleavage motif is shown within an open box followed by a TGF-β superfamily active domain located at the carboxyl end highlighted in grey. Nine cysteine residues found at invariant positions in TGF-β active domains are circled.

**Fig 7:**
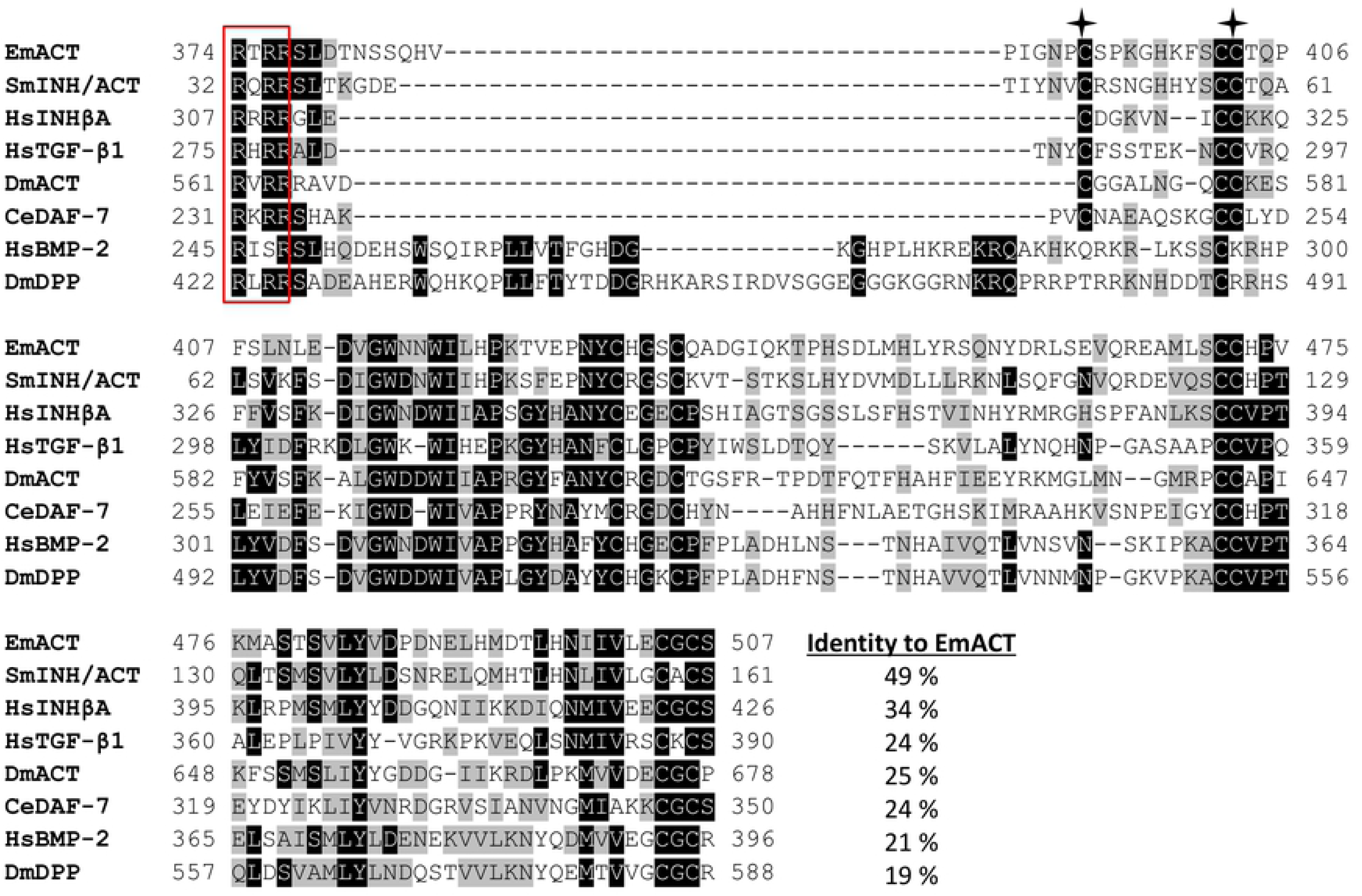
Alignment of the C-terminal amino acid sequences of EmACT and seven other representatives of the TGF-β superfamily. The paired dibasic cleavage motif is shown within a red open box. Residues that are identical are highlighted in black, similarities in grey. Gaps introduced to maximize the alignment are represented by dashes. Two conserved cysteines found only in TGF-β /activin subfamily are shown with asterisks. Numbers at the start and finish of each line correspond to the amino acid numbers in each respective sequence. Accession numbers for the sequences shown are listed in S2 Tab.

Sequence comparisons to several TGF-β superfamily members (Fig 7) and BLASTP analyses of the conserved C-terminal portion of EmACT against protein databases confirmed that its closest relatives are inhibin beta A chains. Highest homologies were detected to SmInAct (49% identical amino acid residues) and human inhibin beta A (34%) (Fig 7). To further confirm that EmACT is an activin/inhibin ortholog, we carried out phylogenetic analyses. The putatively bioactive C-terminal domain of EmACT was aligned to those of several TGF-β superfamily members and the degree of homology was represented on a phylogenetic tree. EmACT clearly clustered with TGF-β/activin subfamily members but not with the BMP subfamily (Fig 8) and, again, showed highest similarity to SmInAct. Taken together, all structural analyses clearly indicated that EmACT is a member of the TGF-β/activin subfamily of TGF-β like cytokines.

**Fig 8:**
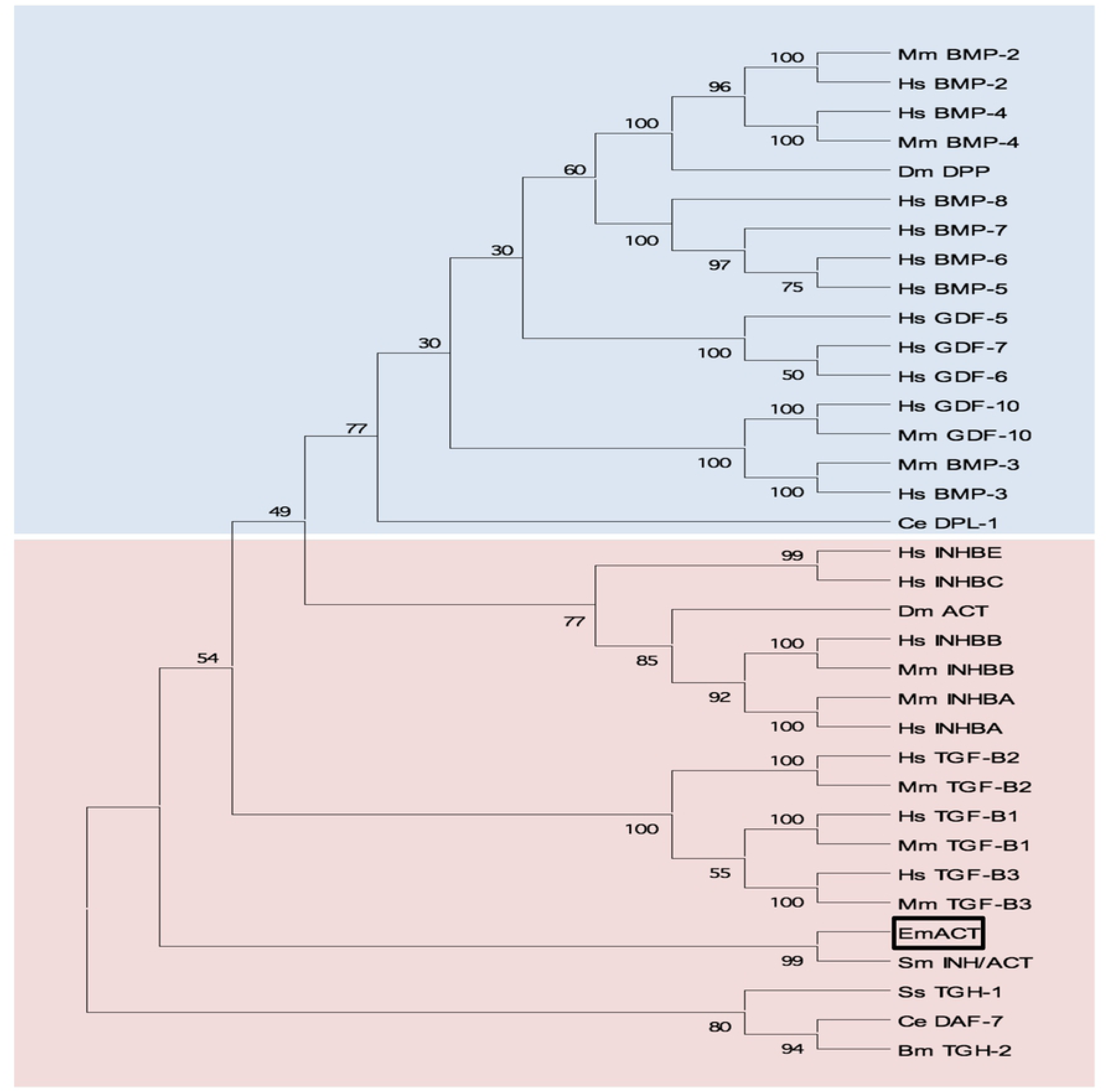
Phylogenetic clustering of EmACT with TGF-β/activin subfamily members. A non-redundant set of TGF-β superfamily members sequences were aligned and an unrooted neighbor-joining tree was computed by MEGA. EmACT is shown clustering with members of the TGF-β/activin subfamily (pink box), but not with members of the BMP/growth differentiation factor subfamily (blue box). Conserved residues in the C-terminal region of each homolog (final 94–106 amino acids) were used in the analysis. Percentages at branch points are based on 1,000 bootstrap runs.

Finally, by using the EmACT sequence as a query in BLASTP analyses against the recently determined genome sequences of other cestodes [32], we identified *Emact* orthologs in *E. granulosus* (EgrG_000178100), *Taenia solium* (TsM_000011500), and *Hymenolepis microstoma* (HmN_000204000), which encoded proteins with 99, 95 and 66% amino acid sequence identity to EmACT, respectively. Hence, the presence of activin A – encoding genes appears to be a common feature of tapeworm genomes.

### Expression of EmACT

Preliminary deep sequencing transcriptome data collected during the *E. multilocularis* whole genome sequencing project [32] indicated that *Emact* is actively transcribed in the *E. multilocularis* metacestode. To closely investigate the formation of the gene product, EmACT, an anti-EmACT antiserum was raised in mice by subcutaneous injection of tag-fused EmACT. This polyclonal serum was used to specifically assess whether EmACT is secreted by *E. multilocularis*. The supernatant of *in vitro* cultivated metacestode vesicles was probed with the anti-EmACT antiserum. We clearly detected reactive proteins of 15- to 25-kDA in the supernatant (Figure 9A), indicating that EmACT has a complex processing and is secreted as different variants of the active protein (∼130 amino acids) by *E. multilocularis* metacestodes.

**Fig 9:**
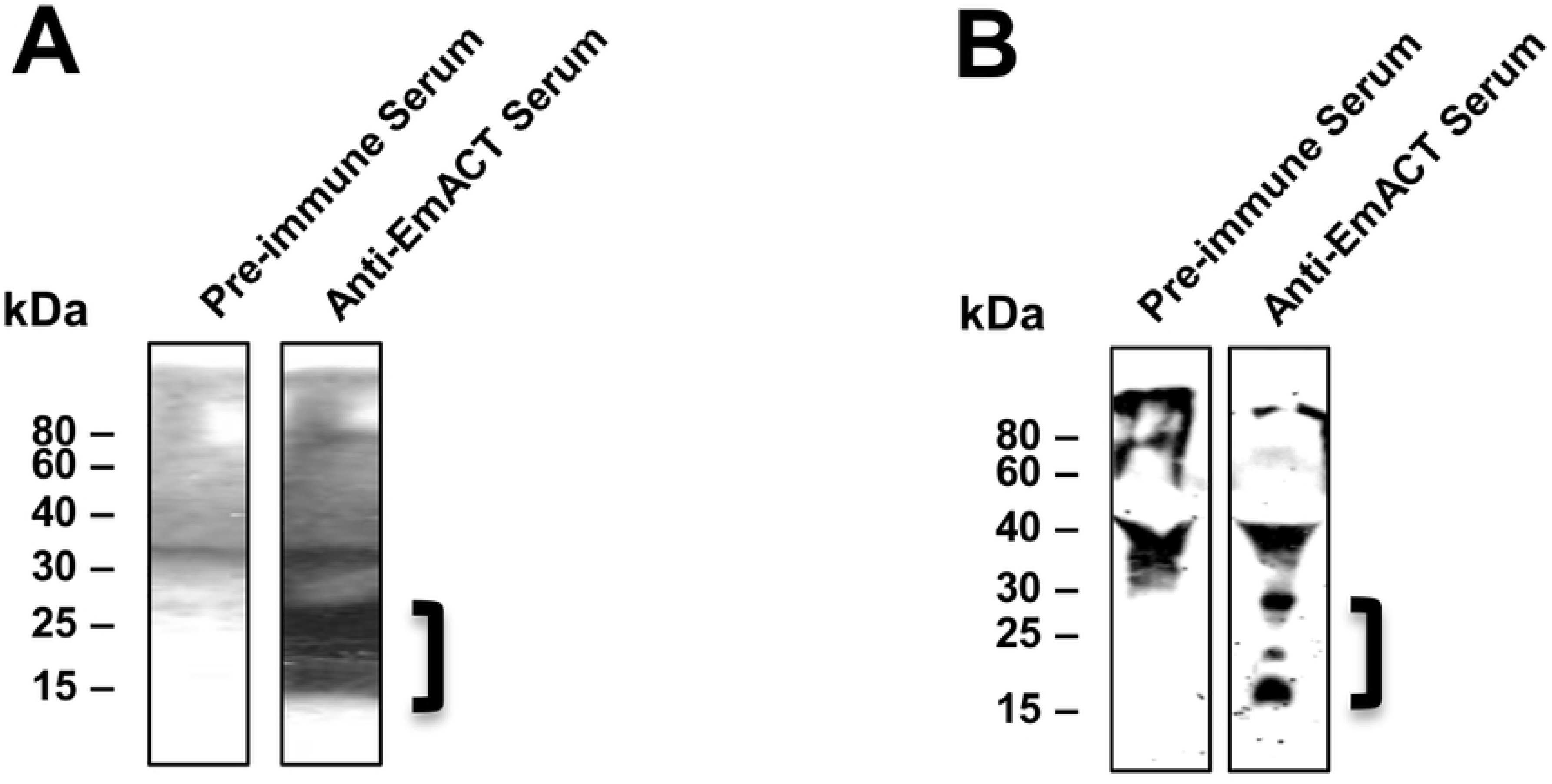
Immunodetection of EmACT. Expression and purification of EmACT fusion protein. EmACT was cloned into the bacterial expression vector pBADThio/TOPO. Competent *E.coli* (Top 10) bacteria were transformed with the Thio-Em*act* plasmid and induced to express the fusion Thio-EmACT protein under arabinose control. A C-terminal histidine repeats fused to the expressed Thio-EmACT fusion protein by the pBADThio/TOPO expression vector was used as target tag for protein purification over Nickel-supplemented beads. Lysates of *E.coli* transformed with pBADThio/TOPO-Em*act* construct before and after arabinose-driven protein expression as well as purified Thio-EmACT were separated by SDS PAGE, blotted over a nitrocellulose membrane and the proteins revealed by Ponceau S staining. The arrow indicates the recombinant Tagged-EmACT. (**A**) Secretion of EmACT by *Echinococcus multilocularis* metacestode vesicles in culture. Shown is a western blotting of ethanol-precipitated MVE/S probed with normal mouse serum or mouse anti-EmACT Immunserum followed by ECL detection and autoradiography. The positions of the molecular mass markers (in kilodaltons) are shown on the left. The bracket indicates the position of EmACT variants. (**B**) Secretion of recombinant EmACT by pSecTag2-*emact*-transfected HEK cells. Shown is a western blotting of the Ethanol-precipitated supernatant of pSecTag2-*emact-* transfected 293T HEK cells probed with either normal mouse serum (or mouse anti-EmACT immune serum followed by ECL detection and autoradiography. The positions of the molecular mass markers (in kilodaltons) are shown on the left. The bracket delimitates the location of recombinant EmACT variants.

For functional characterization of EmACT, the entire protein-coding region was recombinantly expressed in HEK 293 cells under the control of the cytomegalovirus promoter using a mammalian expression system. As shown by Western blotting using the anti-EmACT antiserum, recombinant EmACT (rEmACT) was secreted to the medium by transfected HEK 293 cells as 15- to 25-kDa variants (Fig 9B), which was in agreement with the observed secretion pattern of mature EmACT by the *E. multilocularis* metacestode (Fig 9A). Complementarily, HEK 293 cells were transfected with EmACT containing a Myc tag within the coding sequence, after the furin cleavage site and prior to the mature peptide (S2 Fig) to conceptually enable the secretion of a N-tagged mature EmACT in culture to validate that the assumed complex processing typical of TGF-β superfamily is relevant in EmACT. Indeed, the secretion of c-myc-N-tagged mature EmACT was assessed by immunoprecipitation of the transfected HEK cell supernatant using bead-bound anti-c-myc antibodies and probing the beads’ eluate with anti-c-myc antibodies, revealing specific bands by around 15- to 25-kDa (S2 Fig).

Collectively these results showed that EmACT is secreted by *E. multilocularis* metacestodes as differentially processed variants, which could also be efficiently produced by recombinant expression of EmACT in HEK cells.

### rEmACT induces Treg conversion *in vitro*

Similar to our previous assays using metacestode E/S products, we then investigated whether rEmACT has activin A–like activities. Again, purified naïve CD4^+^CD25^−^ T-cells from spleens and lymph nodes of OT-II.Rag-1-/- mice were isolated and co-cultured with OVA-pulsed DCs. The supernatants of *Emact*-transfected HEK cells (rEmACT) or vector-transfected HEK 293 cells (control) were added to the DC-T-cell co-cultures and the rate of Foxp3^+^ Treg conversion was measured 5 days later by flow cytometry. When compared to the supernatant of vector-transfected HEK 293 cells, rEmACT-containing HEK cell supernatant alone failed to expand Foxp3^+^ Treg but could considerably promote TGF-β-driven Foxp3^+^ Treg conversion (Fig 10). These data indicated that EmACT is unable to induce the *de novo* Foxp3^+^ Treg conversion alone, but synergizes with TGF-β.

**Fig 10:**
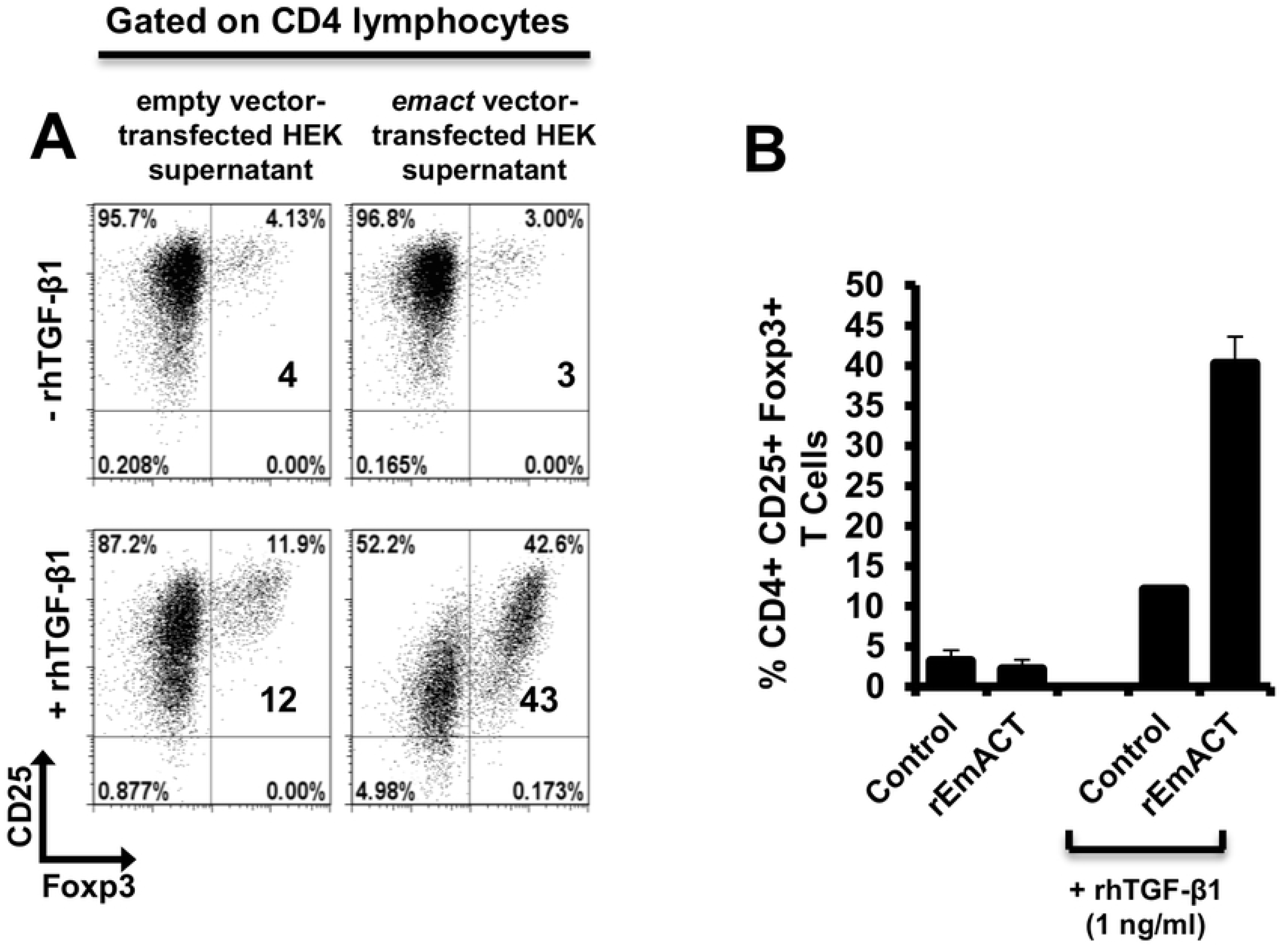
EmACT promotes host TGF-β-dependent Foxp3^+^ Treg conversion in vitro. Freshly generated BMDCs (Day 8) were co-cultured with naïve (CD25^−^) OT-II.RAG-1^-/-^ CD4^+^ T-cells at a DC:T-cell ratio of 1:3 in R10 medium supplemented with OVA peptide (200ng/ml) in the presence of supernatant from pSecTag2-transfected HEK (Control) or pSecTag2-*emact*-transfected HEK (rEmACT) supplemented or not with rhTGF-β1 (1 ng/ml). After 5 days of incubation, cells were harvested and stained for CD4, CD25 and Foxp3 prior to flow cytometry analysis. (**A**) Representative plots of two independently performed Treg conversion assays with supernatant from 2 batches of transfected HEK cells summarized in (**B**). The bars represent the mean ± SD.

### rEmACT induces the secretion of IL-10 by CD4^+^ T-cells *in vitro*

Finally, we also investigated whether rEmACT, like mammalian activin A, is able to stimulate the secretion of IL-10 by CD4^+^ T-cells. Naïve CD4^+^ CD25^−^ T-cells from spleens of C57Bl/6 mice were activated with plate-bound anti-CD3 and anti-CD28 antibodies and the supernatants of *Emact*- or vector-transfected HEK 293 cells were added as test and control, respectively. We noted a considerably higher production of IL-10 in T-cell cultures supplemented with rEmACT-containing HEK supernatant when compared to the control (Fig 11) suggesting that rEmACT can trigger IL-10 release by CD4^+^ T-cells *in vitro*.

**Figu 11:**
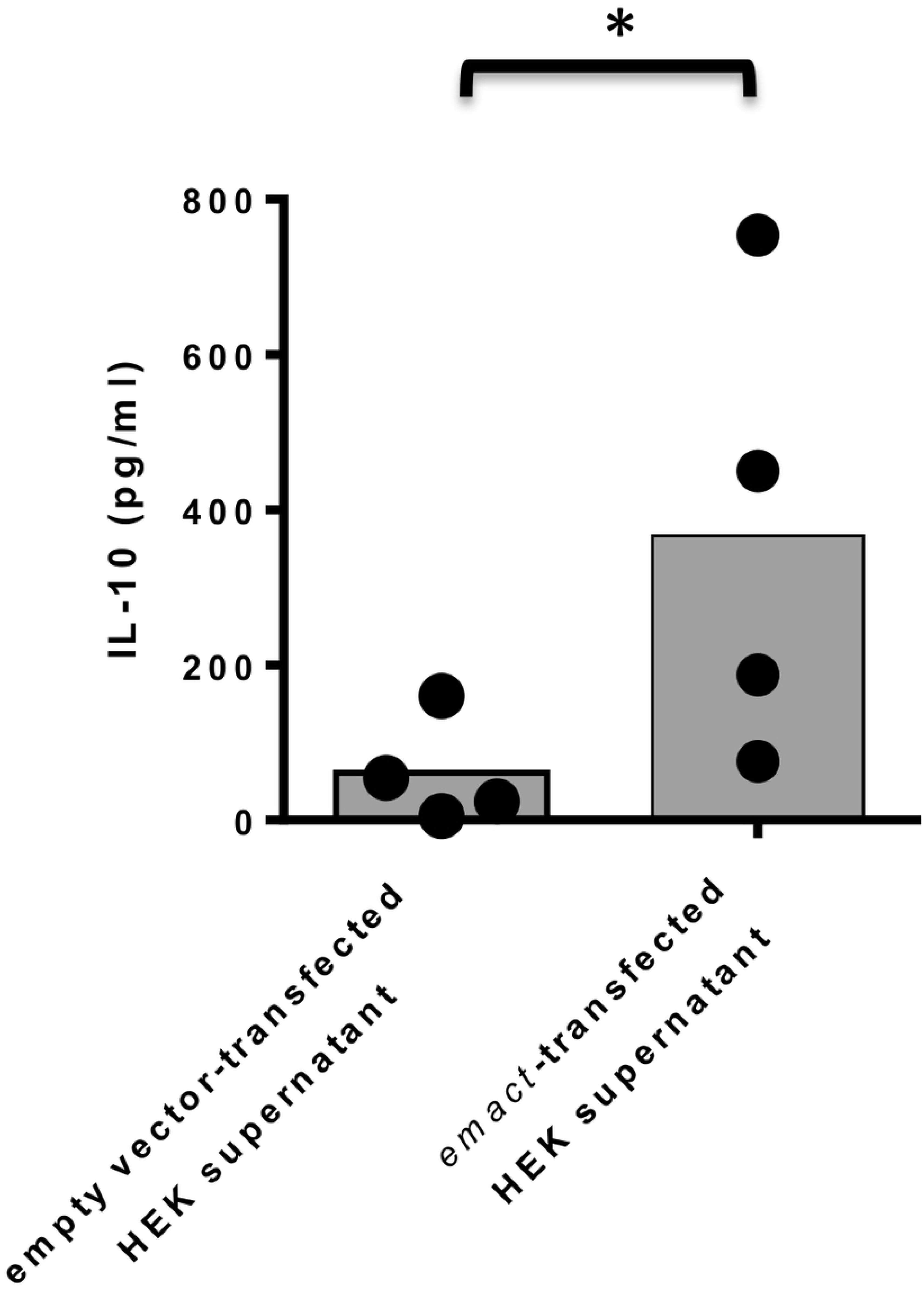
EmACT promotes IL-10 release by CD4^+^ T-cells *in vitro*. CD4^+^CD25^−^T-cells freshly isolated from C57BL/6 mice were stimulated with CD3/CD28 antibodies in the presence of supernatants from pSecTag2-transfected (Control) or pSecTag2-*emact*-transfected HEK cells (rEmACT). After 72 hours, the T-cells supernatants were collected and probed for IL-10 concentration by Elisa. Horizontal bars represent the mean from two independent experiments with T-cells from two different isolations individually activated in the presence of HEK supernatant batches from two different transfections.

## DISCUSSION

AE is a chronic disease characterized by continuous and infiltrative (tumor-like) growth of the *E. multilocularis* metacestode stage over years or even decades within the organs of the intermediate host [1,2,5]. Previous work established that this is associated with considerable immune suppression, provoked by parasite surface structures and metabolites that are released by the actively growing metacestode [1,5,6,8,9,13–17,31,28,37,45]. In other long-lasting helminth infections, regulatory T-cells have been identified as a major contributor to parasite-induced immune suppression [20,46–48] and, very recently, evidence for the expansion of this cell type during secondary AE has been obtained [15,28] with a critical role demonstrated for their immunosuppressive functions in the mitigation of host anti-AE immune response [28]. However, it was not clear from these studies whether Tregs were actively induced by the parasite *in vivo*. Evidence for a certain capacity of *Echinococcus* E/S products to induce Tregs was recently obtained by us in a DC-based Treg expansion assay [6]. This study did not discriminate, however, between mitogenic effects on pre-existing Tregs and *de novo* Treg conversion. Furthermore, the precise mechanism of Treg expansion by *E. multilocularis* remained elusive.

Our present results clearly indicate an active induction of Treg by the parasite as follows: i) Using a well established *in vivo* infection model for secondary AE [8,14–16,37,38,45], we observed a significant expansion of Foxp3+ Treg over effector T-cells in the peritoneum of mice in a time window around 7 days post infection. ii) We demonstrate that Treg formed at this time point are functionally suppressive. iii) Rather than inducing the proliferation of pre-existing Treg, metacestode E/S products can efficiently promote the *de novo* conversion of host Treg *in vitro*, in a TGF-β-dependent manner. iv) In addition to inducing tolerogenic phenotypes in T-cell-priming DC [6], metacestode E/S products can also promote host Treg conversion in a direct, DC-independent manner. Taken together, these data clearly indicate that Foxp3^+^ Tregs can be actively induced by the *E. multilocularis* metacestode to drive host immune suppression during AE.

Interestingly, we also found that metacestode E/S products induce IL-10 release by CD4^+^ T-cells. Whether IL-10 in our assays is produced by Foxp3^−^ or Foxp3^+^ CD4^+^ T-cells is not fully clear at this point. However, since the CD4^+^ T-cell-dependent production of IL-10 in response to parasite E/S products clearly preceded the expansion of Foxp3^+^ Treg (3 days vs. 5 days), Foxp3^−^ cells are likely to be the main source of CD4^+^ T-cell-derived IL-10 in our assays; this would therefore suggest that the parasite-driven Treg expansion and the induction of IL-10 producing T-cells occur largely independently from each other during AE. This is consistent with previous studies on products of the related trematodes *Schistosoma mansoni* [49] and *Fasciola hepatica* [50] for which also Treg expansion and elevated CD4^+^ T-cell dependent IL-10 production had been observed. Hence, such an independent induction of IL-10 producing T-cells would add to the immunosuppression by Foxp3+ Treg and could further contribute to parasite establishment. It also provides for the first time a mechanistic explanation for the elevated IL-10 levels observed in tissues and body fluids of AE patients [9,51,52].

Within the *E. multilocularis* metacestode E/S fraction we identified a component, EmACT, that most likely contributes to the Treg expansion and the induction of IL-10 secretion by T-cells. Like metacestode E/S products, solutions containing recombinantly expressed EmACT promoted Treg conversion *in vitro* and required host TGF-β to do so. Furthermore, rEmACT-containing solutions also triggered the release of IL-10 by host T-cells. We cannot completely rule out at the moment that the metacestode E/S fraction also contains additional factors that could contribute to the observed induction of IL-10 by T-cells and/or to Treg conversion. In fact, this is rather certain given the inability of recombinant EmACT to directly and independently induce the conversion of Foxp3 Treg in our assays. The dependency on host TGF-β suggests an accessory, rather than central, role of this factor in the observed ability of *E. multilocularis* metacestode to expand host Treg. Clearly, other unidentified *E. multilocularis* factors might possess the Treg inducing ability herein reported and in so doing, possibly act in concert with EmACT to promote immunoregulation. In this regard, the *E. multilocularis* genome [32] does, for example, encode homologs of the schistosome ribonuclease omega-1 [53,54] or mammalian BMPs, which have the ability to induce Treg conversion in a TGF-β-dependent manner [55,56]. However, unlike EmACT, these factors have not been reported to induce IL-10 production in T-cells. To further investigate this aspect, we already tried to block EmACT activities in the E/S fraction by using the available anti-EmACT antiserum. Unfortunately, several attempts to immunoprecipitate native EmACT from E/S products using our generated serum failed, indicating that the available antibodies might only recognize the mature protein in its denatured form. To investigate whether additional metacestode E/S components are capable of inducing Treg conversion and/or IL-10 production by T-cells the availability of neutralizing antibodies that recognize native EmACT would thus be necessary. Nevertheless, even if additional parasite components could contribute to the immunosuppressive activities of the metacestode E/S fraction, our experiments on recombinantly expressed EmACT strongly suggest that it is a major component of the cascade of events that promote a Treg and IL-10 rich environment during AE.

In an important previous contribution, Grainger *et al.* [47] demonstrated that E/S products of the nematode *Heligmosomoides polygyrus* can induce Treg *de novo* and suggested a ‘TGF-β mimic’ as the major E/S component to mediate these effects. Although the precise molecule has now been identified in this study as a non-TGF-β superfamily member [57], these authors demonstrated that their molecule acted via the host TGF-β signaling cascade to mediate its effect. In fact, identified nematode TGF-β orthologs also have the capacity to bind to mammalian TGF-β receptors [58]. We now show that a helminth-derived TGF-β-superfamily member can indeed promote Treg (TGF-β dependent) and, at least concerning immune cells, displays clear functional homologies to activin A such as the induction of IL-10 in T-cells [40,41]. Interestingly, our *in silico* analyses could identify similar activin-like molecules in the genomes of other cestodes. Notably, *E. granulosus*, *Taenia solium* and *Hymenolepis sp.* which are pathogens reported to expand Foxp3^+^ Treg and elevated IL-10 production in their hosts [59–63], do all harbor *Emact* orthologs. An implication of this family of molecules in the modulation of the host immune response by these related helminths is therefore possible and merits closer examination.

The fact that E/S products from *E. multilocularis* metacestodes can induce IL-10-secreting and Foxp3^+^ T-cells, which themselves might produce or convey to other immune cells the ability to produce immunosuppressive cytokines like TGF-β and IL-10 [64,65], could explain the high doses of these cytokines found in parasite vicinity during AE infections [10,11,51]. This tightly reconciles with the reported expansion of CD4^+^ Tregs within the periparasitic environment during AE [15,28] and the debilitating role of this cell type on the host ability to control the infection [28]. Since these granuloma also contain CD8^+^ T-cells [10,11,51] we cannot exclude that immunosuppressory mechanisms associated with suppressive CD8^+^ T-cells [15] are also at work. However, since it has been shown that the CD4^+^ fraction is highly important for parasite clearance [38], we think that CD4^+^ Tregs, as expanded by EmACT, are major actors in the impairment of host immunity during AE. Experiments to further verify this have now been performed [28–30] supporting a critical role of this parasite-driven modulation of cell-mediated immunity by Tregs during AE.

Due to the relatively close phylogenetic relationship between helminths and mammalian hosts, it is now clear that they can communicate via evolutionarily conserved signaling systems [66]. Examples are the induction of Epidermal Growth Factor (EGF) signalling in trematodes and cestodes by host derived EGF that binds to evolutionarily conserved EGF receptor kinases [67,68]. We previously demonstrated that also host insulin can stimulate *Echinococcus* development by acting on evolutionarily conserved insulin signaling systems [69]. This apparently also extends to cytokines of the FGF family and the TGF-β/BMP family and respective parasite receptors since host BMP2 has been shown to stimulate a TGF-β family receptor kinase of *E. multilocularis* [71] and similar evidence has also been obtained for schistosomes [72]. It is thus reasonable to assume that parasite-derived cytokines of this family can also functionally interact with TGF-β/BMP receptors of the host. Although we have not yet identified the precise receptor system that is stimulated in T-cells by EmACT, we propose that it acts directly on the Activin receptor-like kinase (Alk) system that is involved in Treg conversion [39,47,73,74]. Further investigations as to which mammalian TGF-β/BMP receptor systems are activated by cestode TGF-β family ligands such as EmACT are clearly necessary.

Although the induction of Treg might be beneficial to *Echinococcus* from the very beginning of the infection, we herein mostly focused on E/S products of the metacestode since we previously showed that E/S products of *Echinococcus* primary cells, which functionally resemble the oncosphere-metacestode transition state [6], did not induce Treg conversion [6] and failed to trigger IL-10 release by T-cells [7]. The reason for these differences might be different composition of the E/S fractions from metacestode vesicles and early primary cells. Indeed in transcriptome data collected during the genome project [32], we already observed clear differences between primary cells and metacestodes in the expression of potentially secreted proteins. Furthermore, we also observed that primary cells secrete a factor EmTIP which induces IFN-γ in T-cells and which is not secreted by the metacestode [7]. Hence, different stages of the parasite (i.e. less protected (primary cells) and well protected (metacestode)) might act differently on T-cells due to a differential E/S profile, and might use different mechanisms for establishing a protective environment. In the case of primary cells, this could include the induction of apoptosis and tolerogenicity in DCs, because they are the first actors at the site of infection [6]. In the case of the metacestode, this could, in addition, involve the formation of Tregs in order to not only contain the host response against the actively growing larva, but most probably also to limit extensive tissue damage in the host.

It has been shown that in addition to immunosuppression, chronic AE is also associated with a Th2 immune response [5]. This could, in part, be a result of a dominant Th2 differentiation of Foxp3^+^ Tregs upon loss of Foxp3 expression observed after the parasite-driven transient expansion of Foxp3^+^ Tregs after 7 days of infection in our assay. This hypothesis is supported by the reported preferential Th2 differentiation of Treg following Foxp3 loss in human T-cells [75]. On the other hand, a more likely contribution of EmACT in the Th2 response reported during chronic AE might come as a result of its conserved functionalities with mammalian activin A which has been shown to promote, in a context-dependent manner, Th2 effector functions [76,77]. Clearly, more experiments are necessary to address these questions.

Taken together, we herein introduce a parasite TGF-β superfamily ligand homologous to activin A, EmACT, which is secreted by the metacestode larva of the tapeworm *E. multilocularis*, and able to promote immunosuppressive features in host T-cells. Moreover, its implicit role in the host immunomodulation by *E. multilocularis* products places EmACT as a therapeutic target for novel anti-*Echinococcus* strategies and a novel tool in the therapeutic regulation of host inflammatory responses.

## ACKNOWLEDGEMENTS

The authors acknowledge Monika Bergmann and Dirk Radloff for technical assistance. We wish to thank Dr. Katrien Pletinckx and Dr. Kerstin Epping for excellent technical advices and support. We also thank Dr. Gabriele Pradel, RWTH Aachen University, Germany for help in the antibody production.

## Supporting information captions

**S1 Tab: List and corresponding accession numbers of gene sequences used**

**S2 Fig: N-term c-Myc tagged EmACT secretion pattern in transfected HEK cells.** The *Emact*-Psectag2 vector construct was modified by site-directed mutagenesis to incorporate a c-Myc tag N-terminal of the EmACT mature peptide sequence and after the furin consensus cleavage motif RTRR. HEK 293 cells were transfected with this construct and kept in culture for collection of supernatant over time (72H). The collected supernatant was processed using the c-Myc tagged protein MILD PURIFICATION KIT Ver.2 (MBL) as per the manufacturer instructions. Briefly, the supernatant was supplemented with anti-c-myc beads for capture of c-myc EmACT mature protein. The incubated beads were eluted with c-myc-containing solutions and the eluate pobed with anti-c-myc for myc-tagged proteins.

